# Feeding habits and the influence of pellet diameter on the feeding responses of the flathead grey mullet (*Mugil cephalus*) in captivity

**DOI:** 10.1101/2021.07.07.451406

**Authors:** Sandra Ramos-Júdez, Neil Duncan

## Abstract

The feeding habits and effect of the diameter of pelleted feeds on the feeding responses of wild juvenile and adult flathead grey mullet (*Mugil cephalus*) in captivity were examined. Optimal pellet size for feeding was defined according to the behavioural responses and ingestion of pellets with different diameters (2, 4, 6, 8 mm) that were dropped into the tank in a random sequence. Larger pellets (6 and 8 mm) were more attractive (lower reaction time, high percentage of capture), but the small to medium-sized pellets (2 and 4 mm) were consumed the most. The optimal size was the 2- and 4- mm pellet diameter for juvenile individuals (365.50 ± 36.90 g; 28.8 ± 0.84 cm) and the 4-mm diameter pellet for adults (937.49 ± 146.54 g; 40 ± 1.12 cm). The preferred feeding area of adult mullet was also studied to estimate preference in relation to pellet characteristics such as floating or sink. Two pellet types, floating or sinking, were offered simultaneously in the water column: at the surface, mid-water column and bottom of the tank. The flathead grey mullet had a preference to feed in the mid-water column and the bottom of the tanks indicating that sinking or slow-sinking pellets would be the optimal feed type in relation to mullet feeding behaviour.

## 1. Introduction

Mullet species are of great ecological and economic importance for both fisheries and aquaculture around the world, which is mainly concentrated in the Mediterranean, Black Sea region and South East Asia (Crosetti, 2016). There is increasing interest in the intensive culture of mullets as it is necessary to diversity marine aquaculture products away from carnivorous fish, to supply demand in local market where mullet is appreciated and to reduce fishing pressure on mullet species. Mullets present good biological characteristics and potential for aquaculture. Positive aquaculture characteristics include, the euryhaline capabilities of many mullets, the fast growth (~ 1 kg per year) (FAO, 2019), the high efficiency in converting food to body mass (EUMOFA, 2020) and the omnivorous diet (Cardona, 2016). The euryhaline capabilities enable that flathead grey mullets (Mugil cephalus) are cultured in freshwater, brackish water, and marine water aquaculture facilities and the omnivorous diet indicates the species’ high potential for culture with diets that do not contain fishmeal and fish-oil. In the Mediterranean region, in addition to flathead grey mullet, other species being cultured are: thicklip grey mullet (*Chelon labrosus*), golden grey mullet (*Liza aurata*), thinlip mullet (*Liza ramada*), and leaping mullet (*Liza saliens*) (Crosetti, 2016).

The expansion of flathead grey mullet aquaculture depends on the development of intensive-farming techniques for which, current limitations include diet formulation, characteristics such as pellet size and feeding practices, i.e., how, when and where to deliver feed to optimize feeding efficiency. It is important to develop feeds that are readily accepted to meet fish nutritional requirements for maintenance, growth and guarantee the developmental competence of breeders and offspring. The flathead grey mullet and many species from the Mugilidae family have been described as omnivorous (see review by Cardona, 2016; Jamabo and Maduako, 2015). This omnivorous classification has focused research on the growth performance of fry and juveniles using feeds formulated with vegetable protein sources (Gisbert et al., 2016; Koven et al., 2020; Torfi et al., 2015), which will allow aquaculture to be a sustainable activity reducing and/or eliminating fishmeal and fish oil from the diets. Despite of the recent interest in Mugilidae diets, the feeding responses of mullets concerning pellet characteristics have received no attention in comparison to other cultured species. As a consequence, there is a lack of commercial pelleted feed for flathead grey mullet juveniles or breeders which, additionally, seem to reject commercially produced pellets during acclimatisation once captured from the wild. Efforts to maximise feed utilisation in mullet species fed artificial feeds in intensive aquaculture should take into account the species feeding behaviour and the characteristics of the pellet.

The physical characteristics of pellets, such as size (shape, diameter and length), colour (contrast), texture (hardness) and density (sinking rate; floating, slow sinking, or fast sinking) that offers a different distribution of the feed into the water, determine the ability of fish to detect the pellet, capture the pellet, and once captured, the acceptance to ingest the feed (Stradmeyer, 1989). Fish species could be reluctant to eat pellets with characteristics that make them not perceived as a desirable food item (Stradmeyer, 1989) and do not match with the type of feeding habits the species occupies; surface, surface/column, column feeder or benthic/bottom feeder (Jobling et al., 2001; Rahman et al., 1992). For instance, catfish, salmon, and shrimp require floating, slow-sinking, or fast-sinking feeds, respectively (Lall and Tibbetts, 2009). In addition, pellet characteristics such as size can directly influence growth, as described for Atlantic salmon (Salmo salar) (Wańkowski and Thorpe, 1979) and Nile tilapia (Oreochromis niloticus) (Azaza et al., 2010). Therefore, providing an adequate pellet particle presented according to the species feeding habits will improve the feed acceptance, growth and development of flathead grey mullet in culture.

The present study aimed to identify pellet characteristics to improve delivery of diets to flathead grey mullet by determining the appropriate pellet size accepted by juveniles and adults that meet feeding habits in intensive captive conditions.

## 2. Methods

### 2.1 Experimental animals

Wild flathead grey mullets were captured from Ebro River and brought to IRTA facilities (Sant Carles de la Ràpita, Spain) and held for 16-17 months before the experiments. Fish were stocked in 10 m^3^ tanks. During the first year, the fish did not accept pellets and were fed at 1.5% of the body weight a soft mixture of 20% sardines, 20% hake, 15% mussels, 10% squid, 10% shrimp and 25% of a ground commercial diet (Skretting, Spain) with 0.1% spirulina. After one year and before the present experiment, fish were presented different sizes of pellet from a commercial on-growing diet for sea bream (*Sparus aurata*) (Skretting, Spain) at 1.5% body weight. Twelve juveniles and twelve breeders were selected to examine pellet size preference and returned back to the main tank. A total of 21 fish were then selected to evaluate the feeding habits of this species. In both tests, fish was allowed to acclimatize to the tanks for 7 days prior to the start of the experiment. Throughout the acclimatization period, fish were fed with polychaetes (TOPSY Bait, Netherlands). Forty-eight hours prior to the behavioural tests the fish were not fed to increase their potential activity and appetite.

The study was performed in accordance with the European Union, Spanish and Catalan legislation for experimental animal protection (European Directive 2010/63/EU of 22nd September on the protection of animals used for scientific purposes; Spanish Royal Decree 53/2013 of February 1st on the protection of animals used for experimentation or other scientific purposes; Boletín Oficial del Estado (BOE), 2013; Catalan Law 5/1995 of June 21th, for protection of animals used for experimentation or other scientific purposes and Catalan Decree 214/1997 of July 30th for the regulation of the use of animals for the experimentation or other scientific purposes).

### 2.2 Preferred pellet diameter

In order to record the response of the flathead grey mullet to pellets of different diameters, three fish were placed per tank in a total of four circular tanks of 2000 L (1.7 m diameter x 1 m depth). The test was performed with twelve juveniles (mean weight: 365.50 ± 36.90 g; mean standard length: 28.80 ± 0.84 cm) and twelve adults (937.49 ± 146.54 g; 40 ± 1.12 cm) separately. In the case of the juveniles, three different diameters (2, 4 and 6 mm) were tested and in the case of the adults, four (2, 4, 6 and 8 mm). The pellets were all the same commercial on-growing diet for sea bream (*Sparus aurata*) (Skretting, Spain).

In each tank, a feeding tube was positioned just below the water surface to introduce the pellets and guide them into the water. The part of the tank that had the feeding tube was curtained with black plastic to screen the fish from any movement or disturbance caused by introducing the pellets. The responses to pellets were recorded with a video system. A video camera (Camera KPC-SN505U, Korea Technology and Communications, Seoul, Korea) encased in a waterproof housing (Integraqua Technology, A Coruña, Spain) was placed inside the water to show a direct view of the entrance of the pellets into the tank and allow observations of the full water column in the tank from surface to bottom (Fig. 1A). The camera was connected to a video recorder (Presentco, Xmotion HD Enterprise 08 Video Recorder, Rister, Barcelona, Spain) and responses were recorded. Pellets were randomly dispensed individually through the tube (143 ± 19 pellets / diameter) at time intervals of 50 s. The test was repeated on three different days at 9:00 h, as the morning was found to be the peak of flathead grey mullet feeding activity, which was diurnal (Collins, 1981).

**Figure 1.**
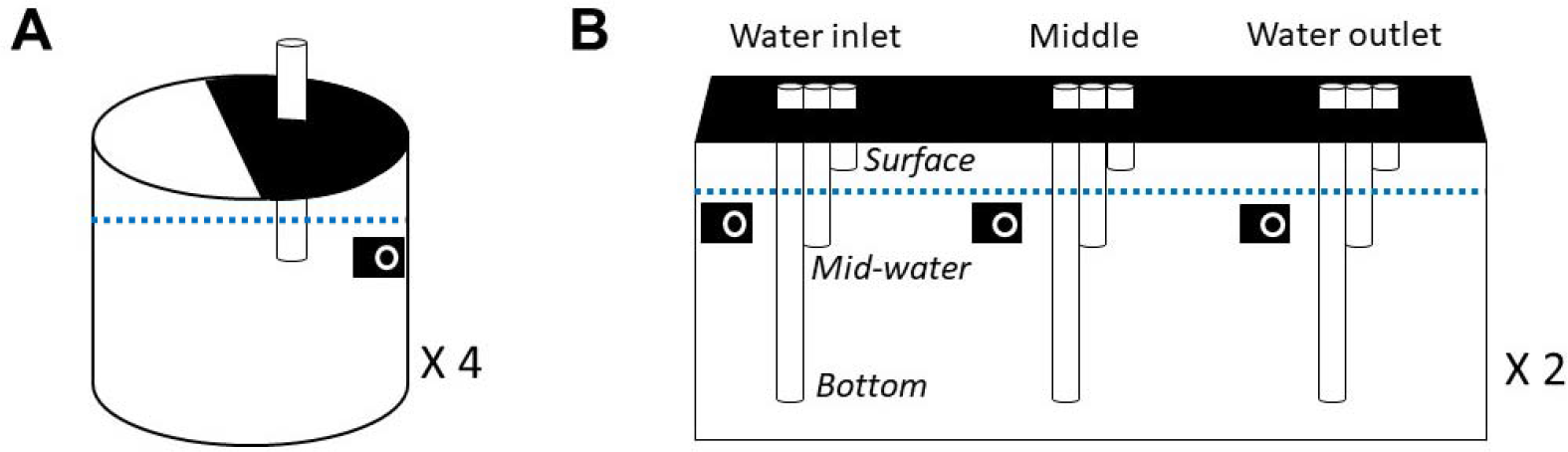
Schematic representation of the tank systems set-up for A) the pellet diameter preference test and B) evaluation of feeding habits. In the first trial, four fiberglass tanks (2000 L each) were prepared with three individuals per tank for adults and juveniles. In the second trial, two 10 m3-tanks were used with 15-16 fish each. Each feeding area (water inlet, middle, water outlet of the tank) was coupled with three tubes with different depths to allow the presentation of food in three levels; surface, mid-water column and bottom. Dotted line indicates water limit. Dark areas represent the part of the tank that was curtained with black plastic to avoid the disturbance of fish caused by the introduction of pellets. Dark square with white circle indicates the camera position.

Different behavioural responses and measures were made from the video recordings. The responses evaluated are based on studies by Stradmeyer (1989) and Smith et al. (1995) and are related to the parameters that influence the final success of feeding; the detectability or attractiveness and acceptability of the food item. Linked to the detectability and attractiveness two parameters were registered, (i) the percentage reaction (rapid movement of the head towards the pellet) or no-reaction of the fish and (ii) the time elapsed since the pellet entered the water and the first catch. Linked to acceptability the following parameters were registered, (i) the percentage of pellets ingested or rejected - those spat out and not recaptured for ingestion- and (ii) the handling time (time elapsed between the first capture and its final ingestion). A preference index (scale from 0 to 1) was calculated dividing the total number of pellets ingested by the number of times the pellet was captured; some pellets were captured more than once as the pellet was spat out and captured a second time (recaptured) or more. The preference index was applied only in those pellets that were ingested. Those pellets with an index close to 0 would be less acceptable than those with an index close to 1. An index of 1 would indicate that all the pellets were ingested when first captured.

### 2.3 Feeding habits

Thirty-one individuals (mean weight: 1044.82 ± 388.92 g; mean standard length: 41.27 ± 3.89 cm) were split in two groups (n = 15, n = 16) and placed in two rectangular 10 m3 tanks of 6.1 × 2.1 × 1.2 m. Three feeding areas were designated in the tanks; water inlet, the middle and water outlet of the tank and a video camera was placed in each feeding area to record the activity as described in the previous trial. In each feeding area, three polyethylene tubes of different lengths were placed to guide pellets into the water column to different depths; surface, in which the tube was placed above the water surface; mid-water column, in which the tube was positioned approximately 25 cm below the water surface to have the pellets fall and appear in the water column; and the bottom, the tube reaching the bottom of the tank (see schematic representation in Fig. 1B). The part of the tank that had the opening of the feeding tubes was curtained with black plastic to avoid disturbance of fish by the introduction of pellets. The tank was illuminated with a strip of LEDs that were positioned in the middle of the tank from the inlet to the outlet and programmed to switch on and off at sunrise and sunset with a gradual increase in intensity. The LEDs were set to emit blue light and the intensity at the water surface increased from 10 lux at the inlet and outlet to 30 lux in the central area of the tank. The lux at the feeding points was approximately, inlet 25 lux, middle 30 lux and outlet 25 lux.

Pellets with the same nutritional formula and the same commercial dietary presentation (Optiline, Skretting, Spain) fabricated for rainbow trout (*Oncorhynchus mykiss*), but with different characteristics were used depending on the depth of the tube in the water column. Floating 4-mm pellets (Optiline AE, Skretting, Spain) were administered through the tube at the surface, and non-floating 4-mm pellets (Optiline AE Ouro, Skretting, Spain) were administered through the tubes positioned mid-water column and the bottom. Five minutes before the feeding activity, a vibrating alarm was presented in both tanks. Subsequently, pellets were administered in the three feeding areas (inlet, middle and outlet of the tank) in one different tube depth (surface, mid-water column or bottom) per feeding area. In this way, the presentation of pellets at three depths were made at the same time with each depth in a different feeding area. Pellets were dropped into the tubes in the following sequence: bottom – mid-water column – surface, in order that pellets were presented to the fish at the same moment. The same pattern of administration (depth and area) was followed for seven days and on the eighth day, it was randomly changed. This sequence was repeated four times over 28 days. This set up ensured that each feeding area in both tanks had periods when feed was administered at each water column depth.

The distribution of feeding fish in the tank was determined by quantifying the number of fish feeding at each feeding point where feed was introduced. The feeding points were a combination of area (inlet, middle and outlet of the tank) and depth at which the pellets were presented (surface, mid-water column and bottom of the tank). The feeding activity was considered to last a total of 25 s, as this was the time that the pellets in the water column had not yet reached the bottom and during which most of the pellets were eaten. Each day, the number of individuals feeding at each feeding point were counted each 5 s during the 25 s feeding period. The proportion of fish feeding at each feeding point was calculated by dividing the number of fish feeding at a feeding point by the total number of fish feeding at the three feeding points on the specific day.

### 2.4 Statistical analysis

The differences between responses to different pellet diameters were evaluated by means of a two-way repeated measures ANOVA with normal distributed data (Shapiro-Wilk test) and equal variance (Levene test), considering the tanks as subjects and the diameter and the days on which the test was performed as factors. Pairwise statistical analysis was performed with the Holm-Sidak post hoc test. The Friedman non-parametric test followed by Wilcoxon test with the Bonferroni correction was applied when the data did not pass the normality test. Feeding habits data did not met normality assumptions. Therefore, the non-parametric Scheirer-Ray-Hare test was applied in each tank. The test is an extension of the Kruskal-Wallis test and represents the alternative to two-way ANOVA (Sokal and Rohlf, 1995). Proportion of feeding fish was the dependent variable and the independent variables were feeding area (inlet, middle and outlet) and depth (surface, mid-column and outlet). The Scheirer-Ray-Hare test was done by applying a two-way ANOVA on ranked data. The H statistic was computed by dividing the Sum of Squares (SS) for each factor and interaction by the total Mean Square (MS). The significance of H was tested as a chi-square variable with the degrees of freedom of the SS being tested. The Dunn’s post hoc test of pairwise multiple comparisons based on rank sums was performed.

A P value of < 0.05 was set to indicate significant differences. Data is presented as mean ± standard deviation (SD) if not indicated otherwise. Statistical analyses were performed using SigmaPlot version 12.0 (Systat Software Inc., Richmond, CA, USA) and MS Excel was used to calculate the H statistic and the significance of H.

## 3.0 Results

### 3.1 Pellet diameter preference

There was no significant effect of the day in which the test was performed on the reaction and ingestion percentages of different pellet diameter in adults or juvenile fish. However, the day in which the test was performed had a significant effect on the manipulation time of pellets in the adults (P = 0.001), as a quicker ingestion was showed in the first day, and on the response time in the case of juveniles (P = 0.005) with a slower response on the first and second days.

The diameter of the pellets had an influence on the reaction of the adult individuals (P < 0.001). The adult flathead grey mullet detected the larger pellets more easily (P = 0.002) (Fig. 2A), although the response time did not significantly differ between the diameters presented (9.7 ± 2.1 s on average) (Table 1). Nevertheless, the initially most attractive sizes were not those consumed once captured, since the diameter was critical in their ability to consume the pellets (P < 0.001). The 4 mm pellets were consumed in a higher proportion followed by the 2- and 6-mm pellets (Fig 2A). The decrease in the preference index from the 2, 4, 6 to the 8-mm pellets (Table 1) indicated that smaller pellets tended to be rejected least often. The preference index’s value for the 8 mm pellets was 0.07 as just one pellet was ingested and this pellet was captured several times before being swallowed. The handling time of the pellets before ingestion was significantly different depending on the pellet diameter (P < 0.001). The smaller the pellet, the shorter the observed manipulation time. Pellets of 2 mm required 2.5 ± 2.4 s to be ingested, 4-mm pellets required 18.7 ± 4.8 s, 6-mm were eaten in 36.5 ± 9.9 and the only 8-mm pellet consumed was consumed after 100 s of manipulation (Table 1).

**Table 1.**
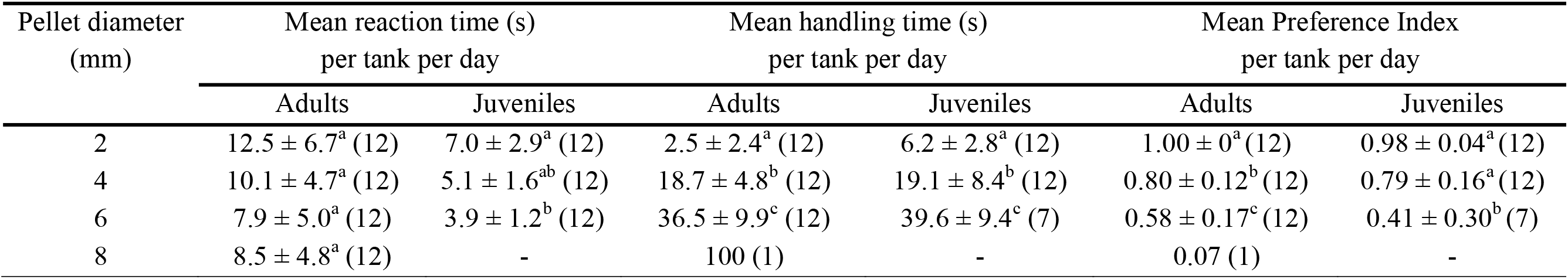
Reaction and handling time (s) of different pellets diameter and Index of preference for adult and juvenile flathead grey mullet (*Mugil cephalus*). Reaction time refers to the time that fish detects the pellet; handling time refers to the total manipulation time by the fish from the first capture of the pellet until ingestion. The preference index was calculated dividing the total number of pellets ingested by the number of times the pellet was captured (scale from 0 to 1, from less acceptable to direct ingestion). Different letters indicate significant differences (P < 0.05). N (in brackets) represents the number of tanks and days in which data was available following the examination of the responses to 143 ± 19 pellets introduced per each diameter in adults and juveniles.

| Pellet diameter<br>(mm) | Mean reaction time (s)<br>per tank per day |  | Mean handling time (s)<br>per tank per day |  | Mean Preference Index<br>per tank per day |  |
| --- | --- | --- | --- | --- | --- | --- |
|  | Adults | Juveniles | Adults | Juveniles | Adults | Juveniles |
| 2 | $12.5 \pm 6.7^a$ (12) | $7.0 \pm 2.9^a$ (12) | $2.5 \pm 2.4^a$ (12) | $6.2 \pm 2.8^a$ (12) | $1.00 \pm 0^a$ (12) | $0.98 \pm 0.04^a$ (12) |
| 4 | $10.1 \pm 4.7^a$ (12) | $5.1 \pm 1.6^{ab}$ (12) | $18.7 \pm 4.8^b$ (12) | $19.1 \pm 8.4^b$ (12) | $0.80 \pm 0.12^b$ (12) | $0.79 \pm 0.16^a$ (12) |
| 6 | $7.9 \pm 5.0^a$ (12) | $3.9 \pm 1.2^b$ (12) | $36.5 \pm 9.9^c$ (12) | $39.6 \pm 9.4^c$ (7) | $0.58 \pm 0.17^c$ (12) | $0.41 \pm 0.30^b$ (7) |
| 8 | $8.5 \pm 4.8^a$ (12) | - | 100 (1) | - | 0.07 (1) | - |

**Figure 2.**
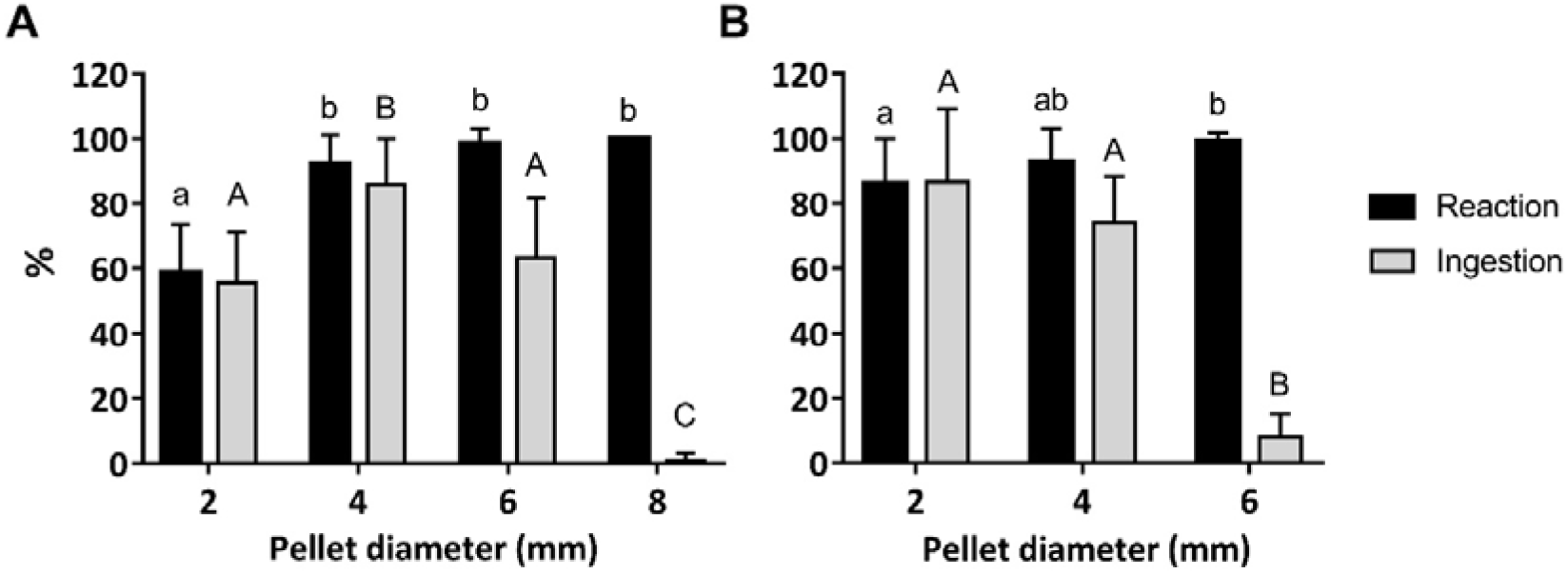
Percentage reaction to pellets and percentage ingestion of pellets (mean ± SD) of different diameters by (A) adult and (B) juvenile flathead grey mullet (*Mugil cephalus*). Three flathead grey mullets were held in each of four tanks that were tested on three days and a percentage reaction / ingestion to 143 ± 19 pellets / diameter was calculated for each tank on each day (n = 12). Different lowercase letters indicate significant differences (P < 0.05) in reaction and capital letters in ingestion. Analysis was performed by the Holm-Sidak post-hoc test after two-way RM ANOVA or the Wilcoxon test with the Bonferroni correction after the Friedman test in no normally-distributed data.

In juvenile individuals, significant differences were found amongst the percentage reaction to 2-, 4- and 6-mm pellet diameters (P = 0.01). Although the juvenile mullet exhibited a high percentage of detection of the three pellet sizes, the reaction to 2 mm pellets was lower than 6 mm pellets (Fig. 2B). Flathead grey mullet also took a significantly longer time to respond to the smaller pellet diameters (Table 1). The time of response was significantly different (P = 0.01) between the biggest (6 mm, 3.9 ± 1.2 s of response) and smallest pellets (2 mm, 7.0 ± 2.9 s), while there was no significant difference between intermediate pellets and biggest. On the other hand, smallest and intermediate pellets (2 mm and 4 mm) were consumed at a significantly higher proportion (Fig. 2B) and their preference index values were closer to 1 (Table 1). The largest diameter (6 mm) was more likely to be rejected. The mean time for pellets to be eaten varied significantly with the diameter (P < 0.01). Ingestion of the pellets took significantly longer with increasing pellet diameter, from 6.2 ± 2.8 s, 19.1 ± 8.4 s to 39.6 ± 9.4 for 2-, 4- and 6-mm pellet diameter, respectively.

### 3.2 Feeding habits

The distribution of fish during the feeding activity significantly depended on the feeding area (inlet, middle and outlet of the tank) (P < 0.001) and the depth in the water column where the food was presented (surface, mid-water column and bottom) (P = 0.009) in both tanks. There was no significant interaction between both factors indicating that the presence of mullet feeding at the surface, in mid-water column or at the bottom did not depend on the feeding area; the inlet, middle or outlet. A significantly higher proportion of fish fed in the middle of the tanks (64 ± 4% and 63 ± 6% in tanks 1 and 2, respectively) (mean ± SEM) in comparison with the inlet (15 ± 2% and 29 ± 6%) and outlet (21 ± 3% and 4 ± 2%) (P < 0.001) (Fig 3A). Regarding the distribution of fish in the water column, tank 1 showed a significantly higher proportion of fish exhibiting feeding activity in the mid-water column (52 ± 5%) (P < 0.001), and tank 2, both in the mid-water column (36 ± 6%) and in the bottom (47 ± 8%) (P < 0.001). Significant lower proportion of fish was found feeding in the water surface (20 ± 3% and 13 ± 4% in tanks 1 and 2, respectively) in comparison with the depth where the fish fed the most in each tank (Fig 3B).

**Figure 3.**
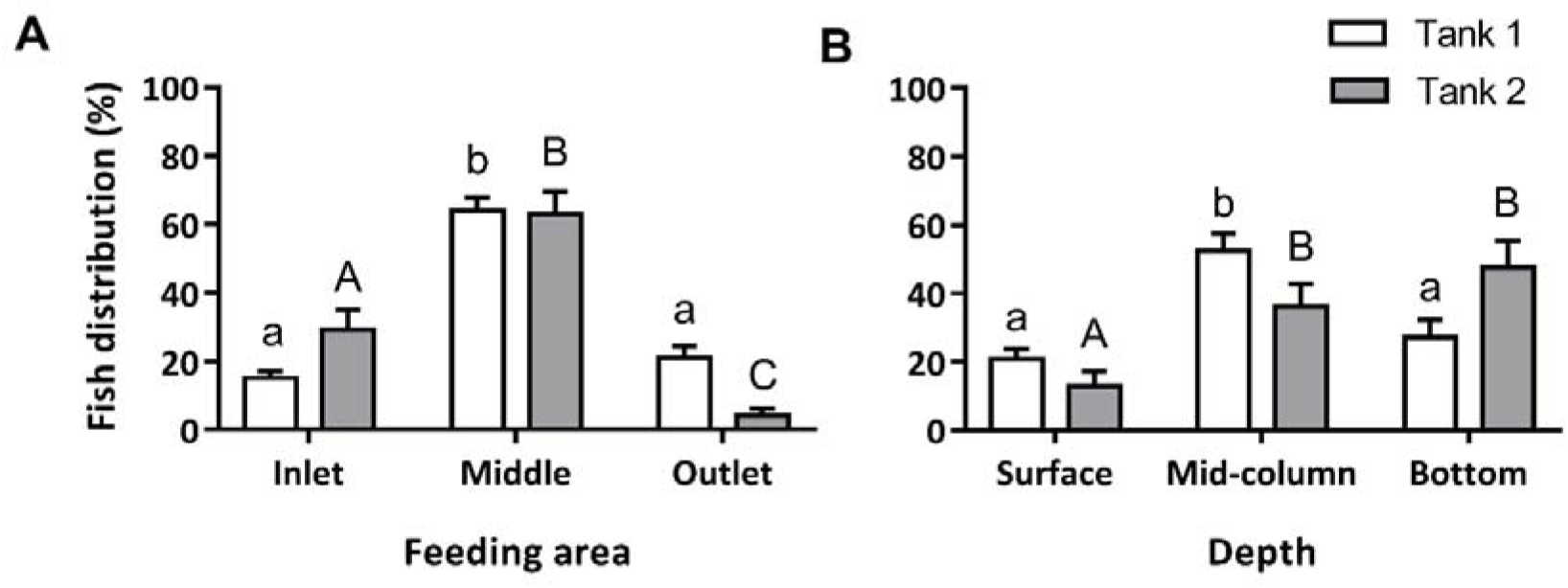
Proportion of fish (mean ± SEM) observed to be feeding in (A) each feeding area (water inlet, middle or water outlet) and in (B) each depth where the food was presented (surface, mid-water column or bottom). Analysis was performed with the non-parametric Scheirer-Ray-Hare with feeding area and depth as factors followed by a Dunn’s post-hoc test. Different small letters represent significant differences (P < 0.05) between feeding areas and depths in tank 1, and different capital letters represent significant differences in tank 2. N per each feeding area and depth is 28 days.

## 4.0 Discussion

The feeding responses of flathead grey mullet recorded in this study reveals the effects of the pellet diameter on the feeding activity and manifests a pattern of feeding habit preferences for this species. The tests were performed in groups to favor responsiveness of the fish for the feed, as flathead grey mullet appear to have a quicker response to feeding in the presence of a feeding group than when isolated (Olla and Samet, 2011). Isolated individuals could become stressed (Pagnussat et al., 2013) and reduce feed intake (Pankhurst et al., 2008). Pellets with different diameters have a different detectability / attractiveness and acceptability that influence the final success of feeding. The percentage of pellets detected and the response time to the pellet represent measures of the pellet detectability and attractiveness. The percentage of pellets consumed and the number of captures followed by rejection (index preference) are measures of the pellet acceptability. Both juveniles (~360 g) and adults (~930 g) of flathead grey mullet responded with higher percentages of responses and a shorter or quicker response time towards the largest and intermediate pellets compared to the smallest pellets. However, the pellet diameters that appeared to be at first the more attractive, were more frequently rejected and were not ingested immediately once captured. The larger pellets were less often captured and the time from capture to ingestion was longer. It is possible that those larger pellets that were finally ingested had softened because of longer contact with water and some manipulation in the mouth of the mullet. Therefore, the present study suggests that the optimum diameter, indicated by those diameters least often missed or rejected and therefore, optimal for feeding, would be 2 and 4 mm for juveniles of ~360 g and 4 mm for adult breeders weighing ~930 g. These results manifest that flathead grey mullet would require pellets of smaller diameter compared to what is expected in other fish species of the same size. If we compare with the manufacturer recommendations for gilthead seabream (*Sparus aurata*), larger diameters are recommended for individuals of similar sizes; 4 to 6-mm pellet diameter for seabream juveniles from 71 to 500 g of weight and 8 mm pellet diameter for seabream > 500 g (Ballester-Moltó et al., 2016). The ability of the sea bream to ingest pellets of larger diameter could relate to this species well-developed chewing apparatus enabling bream to break the pellet into smaller pieces (Ballester-Moltó et al., 2016). In contrast, the flathead grey mullet has been observed to capture the food and keep it in the oral cavity or spit it out and recapture before the pellet was consumed. Mullet did not chew the food item; therefore, the pellets were not observed to be broken into smaller particles. Besides, the teeth of flathead grey mullets are described to be weak and their mandibles are thought to favor the ingestion of sand/mud and organic matter and to select the particles for their final ingestion (Cardona, 2016). Although flathead grey mullets are mainly detritivores and usually feed on plant materials, algae, dinoflagellates, or diatoms, bigger items such as crustaceans, annelids, fish parts, insect parts have also been identified in their stomach contents (Jamabo and Maduako, 2015). It may be possible that flathead grey mullet could eat bigger diameter pellets, but softer than the ones tested in the present study. Soft pellets of a given diameter have been more acceptable in several species (Knights, 1985) including salmon juveniles (Stradmeyer, 1989). Therefore, it would be interesting to test pellets of different hardness and evaluate the relationship between the hardness of the pellet, the diameter, and acceptability.

In regard to flathead grey mullet feeding habits, the present study identifies that feeding mullet exhibited a distribution throughout the water column, including the water surface and the bottom. Nevertheless, it was detected that mullet have a preference for feeding in the water column and the bottom rather than in the surface. This result matches with the natural feeding habit of the species, which feeds on suspended plant materials and forages the bottom (Cardona, 2016). The same feeding habits were identified in juveniles held in tanks by Jimenez-Rivera et al. (2021), in contrast with Ghion (1986) which identified that although eating in all parts of the tank, the feeding increased when juveniles were fed at the surface. According to the present results, slow-sinking or fast-sinking feeds would meet the preferred feeding habits in the flathead grey mullet. The present study, also detected a preference for feeding location in the tank, the middle of the tank. There are several possible explanations for this preference as the tank environment was not completely uniform. For example, the inlet area had more disturbance from the incoming water and the outlet area may have had slightly lower water quality. In addition, the blue light intensity emitted by the LEDs was slightly higher in the middle of the tank than at the ends (inlet and outlet) and might have affected motivation for feeding in these areas.

## 5.0 Conclusion

The present study has identified the preferred characteristics of pellets in terms of size; diameter (2 and 4 mm for juveniles and 4 mm for adults), and density (sinking or slow-sinking pellets) according to the feeding responses and feeding habits of juveniles and adults of flathead grey mullet (feeding in the water column and the bottom). The present study provides a basis for the future development of an optimal pelleted diet for this species in intensive culture conditions.

## 6.0 Acknowledgements

Thanks to IRTA staff for their technical help, to Alex Rullo Reverté and Mario Villalta Vega for help performing the tests and to Anna Belén Castilló Pérez who helped in video evaluation. This study was funded by the National Institute for Agricultural Research and Technology and Food [grant number FEDER-INIA RTA2014-0048]; the Spanish Ministry of Science and Innovation [grant number RTI2018-094710-R-I00]; and the Seventh Framework Program of the European Community for research, technological development and demonstration actions [grant number KBBE-2013-07 single stage, GA 603121, DIVERSIFY]. The participation of S. Ramos-Júdez was financed by a FI-AGAUR (Generalitat de Catalunya) predoctoral scholarship co-financed by the European Social Fund.

